# c-Jun regulates postpartum β-cell apoptosis and survival downstream of prolactin signaling

**DOI:** 10.1101/2025.02.26.640362

**Authors:** Jin-Yong Chung, Nelmari Ruiz-Otero, Ronadip R. Banerjee

**Author notes:** Corresponding author: Ronadip R. Banerjee.

## Abstract

**Objective:** Pregnancy and postpartum states drive dynamic expansion and regression of maternal β-cell mass. Little is known about what regulates postpartum regression. We recently profiled murine islets at different time points from late gestation to early postpartum to identify regulators of β-cell apoptosis or survival. One hit was c-Jun, a transcription factor which regulates proliferation, apoptosis, and survival in various tissues and cells. Here, we examine c-Jun regulation and function during gestation and postpartum and in murine and human islets.

**Methods:** To examine regulation of c-Jun within β-cells we used a mouse genetic model lacking β-cell prolactin receptor (PRLR) and stimulation of cultured islets with recombinant prolactin. We used chemical inhibitors of signal transduction pathways to examine signaling downstream of PRLR. TUNEL was used to detect endogenous or dexamethasone-induced apoptosis. Q-PCR, Western blotting, and immunostaining were used to assess gene and protein expression. Knockdown of c-Jun in MIN6 cells was accomplished using siRNA and lentiviral-shRNA. We examined both murine and human islets and tissues.

**Results:** We found that expression of c-Jun transcript in murine β-cells is temporally restricted to late gestation and early postpartum and requires prolactin signaling. Moreover, c-Jun protein expression was mutually exclusive with apoptotic β-cells identified by TUNEL staining. In both murine and human islets, prolactin treatment is sufficient to induce c-Jun expression and downstream MAPK/ERK signaling. Pharmacologic inhibition of c-Jun blocks prolactin-mediated survival of β-cells following pro-apoptotic stress, including failure to upregulate pro-survival factors Bcl2l1 (Bcl-xL) and Birc5 (Survivin). Finally, human islets during pregnancy also exhibit increased c-Jun expression in β-cells, but c-Jun induction is absent in β-cells from pregnant donors with gestational diabetes (GDM).

**Conclusions:** c-Jun contributes to pro-survival effects of lactogens downstream of PRLR / MAPK signaling in β-cells. c-Jun regulation is conserved in human islets and pregnancy and dysregulated in GDM.

## 1. Introduction

Expansion and reduction of maternal β-cells during pregnancy and postpartum is a physiologic example of rapid and dynamic changes in β-cell mass occurring outside of organism development [1–3]. Thus, elucidation of molecular mediators governing these processes has the potential to yield new insights into how the amount of functional β-cell mass is controlled. Furthermore, as deficiency of functional β-cells is a key component of diabetes pathogenesis, mechanisms regulating functional β-cell mass has therapeutic potential to impact disease management [4–6].

Regulators of gestational β-cell mass expansion have been identified in rodent models, including through transcriptomic studies [7–11]. However, very few studies have examined how β-cell mass is restored to levels comparable to pre-pregnancy following parturition, also known as postpartum “regression” of β-cell mass [12–14]. The cellular mechanism of regression remains controversial and has been reported to occur via reduction in β-cell size, increased β-cell apoptosis, and macrophage phagocytosis of β-cells [13–15]. Lactogen signaling, which activates PRLR, is most widely known for regulating milk production in the mammary glands but is also a critical regulator of gestational β-cell adaptations [8, 16–18]. Recent studies have revealed that lactogen-signaling, via downstream serotonin production also regulates β-cells in the postpartum and lactating periods [14, 19]. However, a comprehensive examination of signals and mechanisms contributing to postpartum regression has not been performed.

To address this question, we recently used single-cell RNA sequencing to identify transcriptional changes within maternal islets in mice from non-pregnant, pregnant and postpartum states [7]. We found that multiple immediate early genes were induced in β-cells during late pregnancy and early postpartum, including c-Jun.

c-Jun is a transcription factor that is a member of the AP-1 transcription family with established roles in regulating proliferation, apoptosis, and survival in multiple cell types and contexts [20, 21]. However, only a handful of studies have examined c-Jun or AP-1 function in β-cell lines or murine islets, and none have examined c-Jun or other AP-1 components in pregnancy or postpartum adaptations. For example, studies of INS-1 and MIN6 cells stimulated with glucose and cAMP found induction of IEGs including c*-Fos, c-Jun, JunB* and *nur-77* (aka *Nr4a1*) by glucose and GLP-1 cotreatment, and validated direct regulation of only a single downstream target gene, *Srxn1* by c-Fos [22, 23].

Here we examine c-Jun regulation in gestational and postpartum islets, and how it regulates islet responses to pro-apoptotic stress, and find that c-Jun, downstream of PRLR and MAPK signaling directly regulates survival from a pro-apoptotic stressor in murine and human islets. c-Jun pro-survival effects are contributed to by induction of pro-survival genes *Birc5* and *Bcl2l1*.

## RESULTS

### 2.1 Regulation of c-Jun during pregnancy by Prolactin / PRLR signaling

We first sought to identify signals responsible for c-Jun induction in late gestational and postpartum β-cells. Considering the established role of PRLR as a critical regulator of β-cells during pregnancy, we investigated whether c-Jun was regulated by lactogen signaling. Pancreatic islets exhibited strong c-Jun staining in late pregnancy within murine β-cells, which persisted into the early postpartum period (PP5) (**Figure 1A**), consistent with our prior data [7]. By contrast, mice lacking PRLR signaling within β-cells failed to exhibit c-Jun expression in β-cells either in pregnancy or postpartum (**Figure 1A-B**).

**Figure 1.**
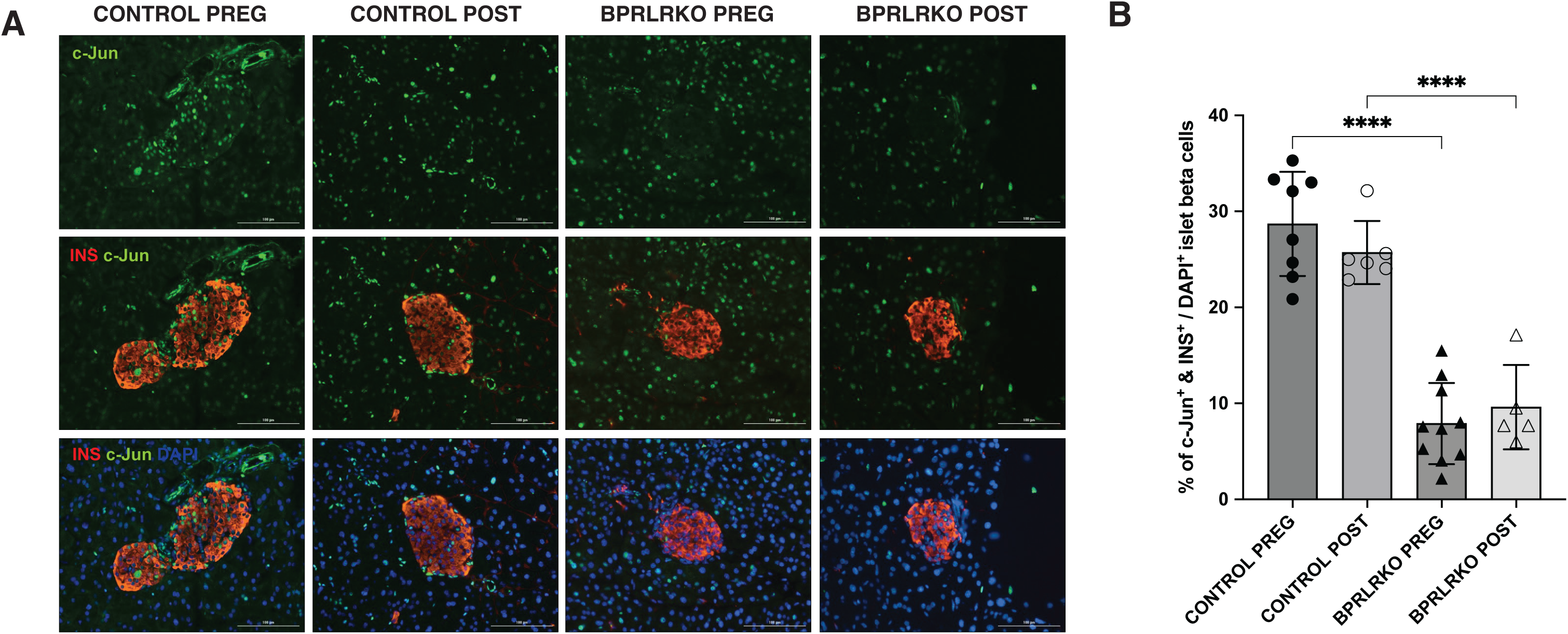
PRLR signaling within β-cells is required for gestational and postpartum c-Jun induction in murine islets. (A) Immunostaining of islets from murine pregnancy (PREG; gestational day 16.5) or postpartum (POST; postpartum day 5) in control (PRLRf/f) or BPRLRKO mice (PRLR^f/f^; RIP-Cre). c-Jun (green) and insulin (red); nuclei were stained with DAPI (blue). (B) Quantification of c-Jun+ cells as a percentage of β-cells (insulin+). Data in B were analyzed using a 2-way ANOVA. Data are represented as mean ± SEM. Statistical significance was defined as ****p ≤ 0.0001.

As c-Jun has been shown to mediate both acute and chronic stress responses in a variety of contexts in various tissues, we next considered whether the increase in c-Jun was specific to pregnancy or might indicate a generalized metabolic stress response. We examined islets from mice fed a high-fat, high-sucrose diet for 24 weeks but unlike islets in pregnancy, c-Jun immunostaining was not detected (**Supplementary Figure S1**). Together, these results indicate upregulation of c-Jun during late pregnancy is not due to islet stress but rather is specifically induced by PRLR signaling during gestation and postpartum.

Having established that PRLR signaling is necessary for c-Jun gestational induction, we next sought to identify whether activation of PRLR signaling was sufficient to induce c-Jun expression. We treated cultured murine or human islets with their species-specific recombinant prolactin (PRL) to stimulate PRLR signaling. PRL increased c-Jun protein levels significantly in both mouse and human islets in culture (**Figure 2A&B**). Co-immunostaining with insulin confirmed increased c-Jun levels were detected in β-cells at 24 hours post-treatment in murine islets (**Figure 2A**). A small number of c-Jun^+^ cells were also observed in non-β-cells. Western Blot quantitation indicated that c-Jun increase was approximately 2-fold at 24 hours (**Supplementary Figure S2A**). Similarly, in islets from a human female donor (see **Table 1** for additional donor information), treatment with recombinant human PRL increased c-Jun^+^ nuclei (**Figure 2B**) and phospho-c-Jun levels (**Supplementary Figure S2B**). As in murine islets, c-Jun immunostaining was predominantly seen in β-cells, but was also observed in some insulin^-^ cells. Together, these data demonstrate that both murine and human islets increase c-Jun within β-cells following PRL treatment, indicating that increased c-Jun following lactogen stimulation is conserved across species.

**Figure 2.**
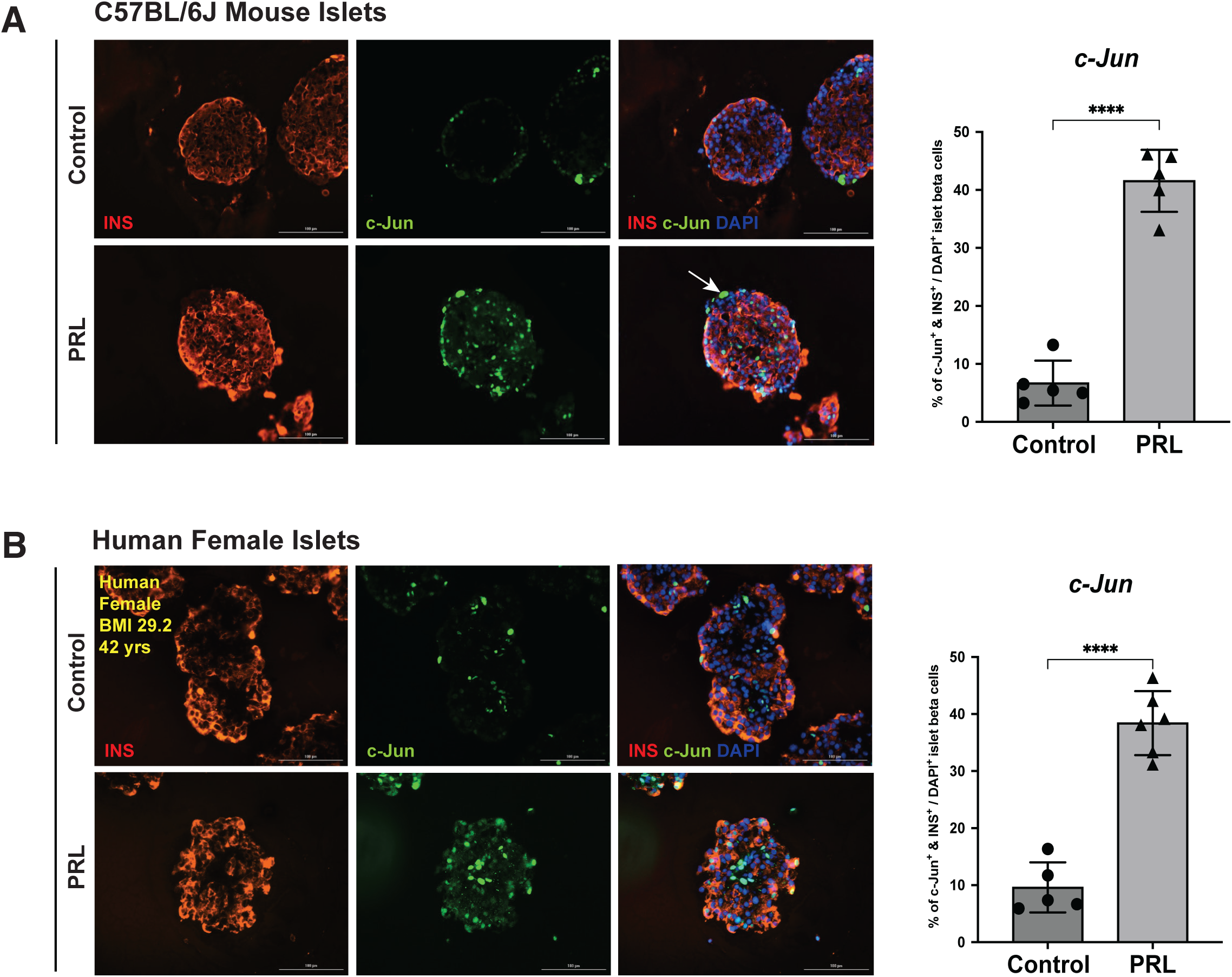
c-Jun is induced in β-cells in murine and human islets cultured with prolactin. Immunostaining of c-Jun in (A) murine and (B) human islets following treatment of cultured islets with recombinant prolactin (PRL) for 24 hours. c-Jun (green) and insulin (red); nuclei were stained with DAPI (blue) (representative images). Graphs show quantification of c-Jun+ cells as a percentage of β-cells (insulin+). Data in A and B were analyzed using a student’s t-test. Data are represented as mean ± SEM. Statistical significance was defined as ****p ≤ 0.0001.

**TABLE 1:** DONOR INFORMATION ON HUMAN ISLETS & TISSUES.

| **DONOR** | **AGE** | **SEX** | **BMI** | **RACE** | **OTHER (pregnancy etc)** |
| --- | --- | --- | --- | --- | --- |
| **nPOD #6401** | 25 | F | 31.3 | Hispanic/Latino | NP#1 |
| **nPOD #6253** | 19 | F | 34.3 | African Am | NP#2 |
| **nPOD #6383** | 22 | F | 31.3 | African Am | PREG#1 (40 wks gestation) |
| **nPOD #6257** | 19 | F | 45.9 | American Indian | PREG#2 (12 wks gestation) |
| **nPOD #6082** | 33 | F | 34.4 | Caucasian | PREG#1, GDM (33 wks gestation) |
| **nPOD #6239** | 35 | F | 50.9 | African Am | PREG#2, GDM (29 wks gestation) |
| **IIDP H. Islets** | 41 | F | 38.4 | American White | ID# RRID:SAMN36823227 |
| **IIDP H. Islets** | 42 | F | 33.3 | Hispanic/Latino | ID# RRID:SAMN40992344 |
| **IIDP H. Islets** | 49 | F | 21.2 | American White | ID# RRID:SAMN39643537 |

### 2.2 MAPK is the signal transduction mediator of c-Jun downstream of PRLR

PRLR signaling regulates intracellular metabolism and transcriptional regulation via multiple signal transduction pathways, including Stat5, MAPK and PI3K/Akt signaling [1, 2]. To clarify the signal transduction mechanisms responsible for c-Jun upregulation following PRLR activation by PRL treatment, we examined these canonical mediators of PRLR signaling. In cultured murine islets, PRL treatment induced robust Stat5 phosphorylation within 30 minutes which peaked at 1 to 3 hours (**Figure 3A**) validating lactogen responsiveness [16, 24]. c-Jun induction was similar, peaking about 3 hours post-stimulation, and remaining significantly above baseline levels at 24 hours (**Figure 3A**). *Jun* mRNA increase was first detected at 1-hour post-stimulation (**Figure 3B**). By contrast, Jun family members *JunB* and *JunD* expression levels were essentially unchanged with PRL stimulation (**Supplementary Figure S3**). Because the increase in c-Jun protein levels (0.5 hours) occurs prior to increased *Jun* gene expression, these results indicate that both post-transcriptional and transcriptional mechanisms contribute to early increases in c-Jun protein levels following PRL stimulation.

**Figure 3.**
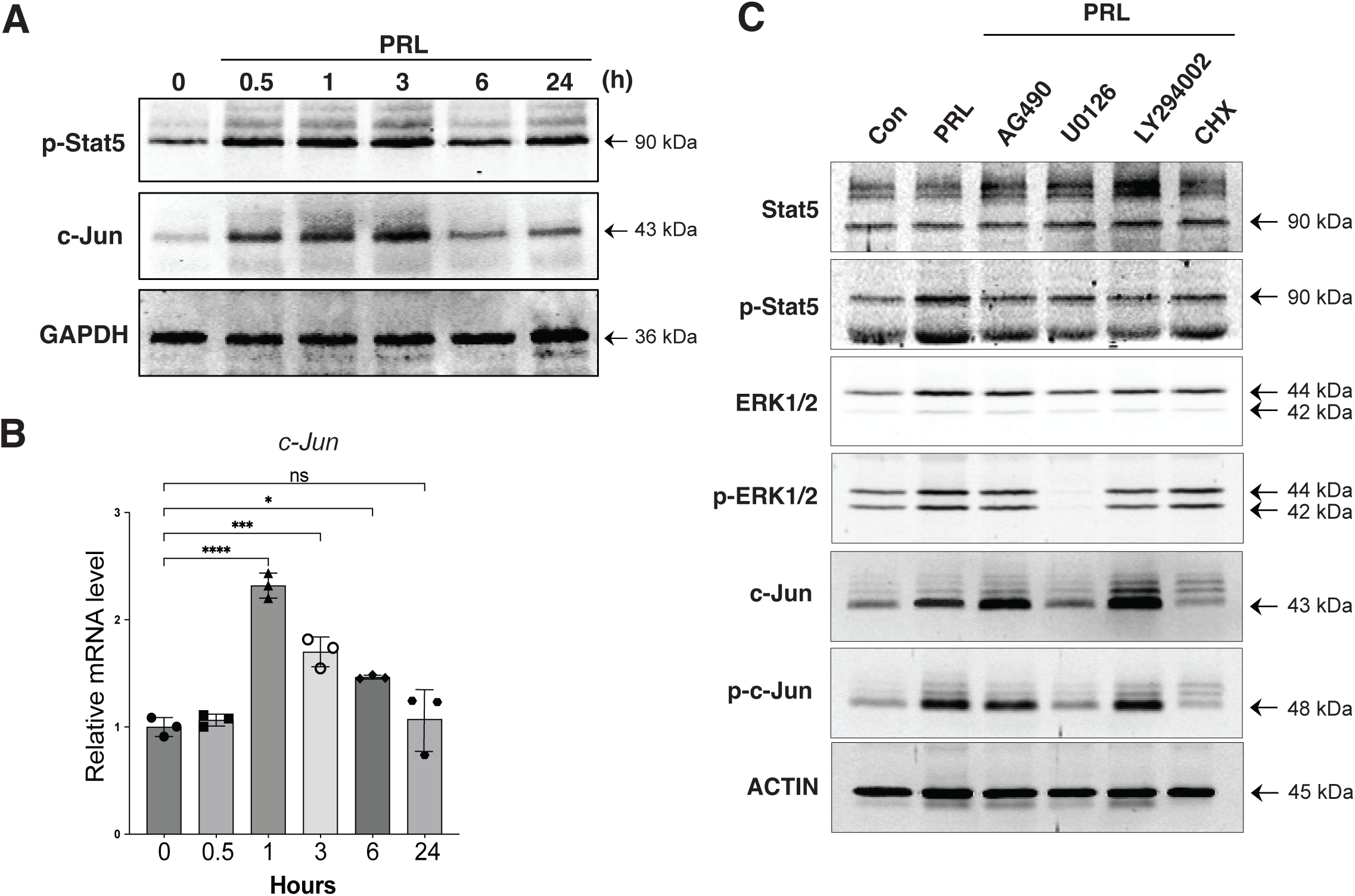
c-Jun is induced by prolactin in islets via prolactin receptor (PRLR) and MAP kinase signaling. (A) Time course of c-Jun protein induction following PRL treatment using Western blot. Phospho-Stat5 (p-Stat5) indicates activation of PRLR activation. GAPDH was used as an internal control. (B) *Jun* mRNA expression by Q-PCR following PRL treatment. Data are represented as mean ± SEM. Statistical significance was defined as *p ≤ 0.05, **p ≤ 0.01, ***p ≤ 0.001, and ****p ≤ 0.0001. (C) Inhibitors of Stat5 (AG490), MAPK (U0126), PI3K (LY294002) and protein synthesis (CHX) indicate that PRLR induces c-Jun via MAPK/ERK signaling.

We next sought to identify the signal transduction mediator(s) critical for c-Jun expression. To do so, we examined 3 hours post-PRL stimulation as the timepoint with maximal c-Jun levels. Inhibition of Stat5 signaling using the inhibitor AG490 blunted P-Stat5 induction, but surprisingly did not block the increase of c-Jun protein following PRL treatment (**Figure 3C**). The PI3K inhibitor LY294002 increased c-Jun levels, consistent with its effects in neuronal cells [25]. By contrast, the MEK1/2 (MAPK) inhibitor U0126, blocked ERK phosphorylation normally induced by PRL, and abrogated the increase in c-Jun and p-c-Jun (**Figure 3C**). Finally, treatment with the protein synthesis inhibitor cycloheximide significantly inhibited the PRL-mediated increase in c-Jun, and p-c-Jun, but did not affect Stat5, ERK1/2, or actin levels, indicating that new protein synthesis is critical for increasing c-Jun following PRL stimulation. In sum, PRLR signaling is critical for c-Jun induction, which occurs via MAPK rather than Stat5, previously established as the predominant gestational mediator of PRLR signaling [26].

### 2.3 c-Jun induction following prolactin treatment of β-cells protects against dexamethasone-induced apoptosis

c-Jun can promote either apoptosis, proliferation, or survival in a cell and context-dependent manner. As the postpartum regression of β-cells occurs shortly following parturition when significant β-cells apoptosis occurs [7, 13], we hypothesized that c-Jun might regulate either β-cell apoptosis or survival. To mechanistically interrogate these possibilities, we used an in vitro system in which β-cell apoptosis was triggered with the glucocorticoid dexamethasone (DEX). DEX was chosen as a physiologically relevant apoptosis inducer because the glucocorticoid surge in late pregnancy and at parturition is thought to be a major mediator of β-cell apoptosis [27, 28]. DEX treatment increased apoptosis of β-cells in a dose-dependent manner as expected (**Supplementary Figure S4**). PRL has been shown to antagonize the pro-apoptotic effects of DEX on β-cells, but the specific mechanism is still incompletely understood [27–30]. In cultured murine islets, control and PRL-treated islets exhibited very low apoptosis rates, whereas DEX treatment significantly increased apoptosis detected as TUNEL^+^ cells (**Figure 4A-B**). PRL treatment caused significant increase in Jun^+^ islet cells. Cotreatment with PRL and DEX increased c-Jun, and significantly reduced TUNEL^+^ cells. However, addition of c-Jun peptide, an inhibitor which blocks c-Jun interaction with JNK (see Methods) blocked the protective effect of PRL against DEX-induced apoptosis (**Figure 4A-B**). Surprisingly, DEX treatment alone increased c-Jun expression, particularly in the islet periphery. However, following DEX treatment, c-Jun^+^ cells and TUNEL^+^ cells were mutually exclusive, suggesting that DEX and PRL increase c-Jun in different cells by different downstream mediators (**Figure 4A-C**). Taken together, these results suggest that the protective effects of PRL against DEX-induced apoptosis are mediated by c-Jun. We next sought to determine whether human β-cells would also be protected against apoptosis by PRL via c-Jun. We obtained human islets from the Integrated Islet Distribution Program (IIDP) from two females of reproductive age (**Table 1**) and treated the islets with DEX, PRL and c-Jun peptide (representative experiment in **Figure 5A**). As with murine islets, PRL treatment induced robust c-Jun expression, but not apoptosis as indicated by TUNEL staining. DEX treatment induced apoptosis, as well as some scattered c-Jun expression, but not within apoptotic cells. Cotreatment with DEX and PRL induced robust c-Jun expression and significantly reduced TUNEL^+^ cells as compared with DEX alone. c-Jun peptide treatment increased TUNEL^+^ cells to levels comparable to DEX treatment alone (**Figure 5A-B**) but did not affect the number of cells expressing c-Jun (**Figure 5C**). Together, these results suggested that in human islets, as with murine islets, c-Jun induction is critical for the protective effect of PRL against DEX-induced apoptosis.

**Figure 4.**
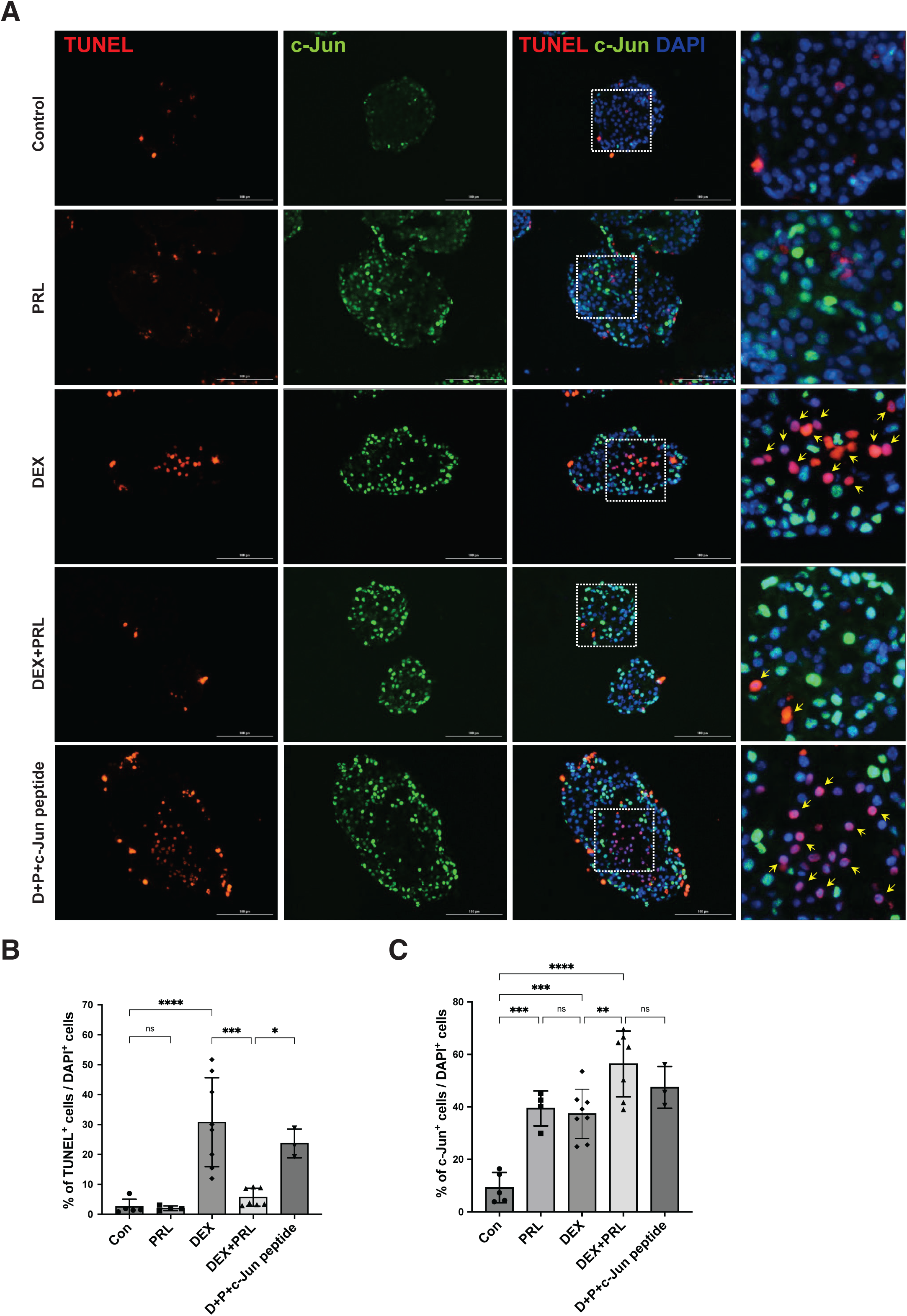
Prolactin-mediated rescue of dexamethasone-induced β-cells apoptosis requires c-Jun activity in mouse islets. (A) Immunostaining of c-Jun (green) following treatment with PRL, dexamethasone (DEX), DEX+PRL, or DEX+PRL+c-Jun peptide (inhibitor). Apoptosis is indicated by TUNEL+ nuclei (red). All nuclei are stained with DAPI (blue). (B) Quantification of percentage of apoptosis (TUNEL+ cells) / total islet cells (DAPI+) (C) Quantification of c-Jun+ cells / total islet cells (DAPI+). Data in B and C were analyzed using a 2-way ANOVA. Data are represented as mean ± SEM. Statistical significance was defined as *p ≤ 0.05, **p ≤ 0.01, ***p ≤ 0.001, and ****p ≤ 0.0001.

**Figure 5.**
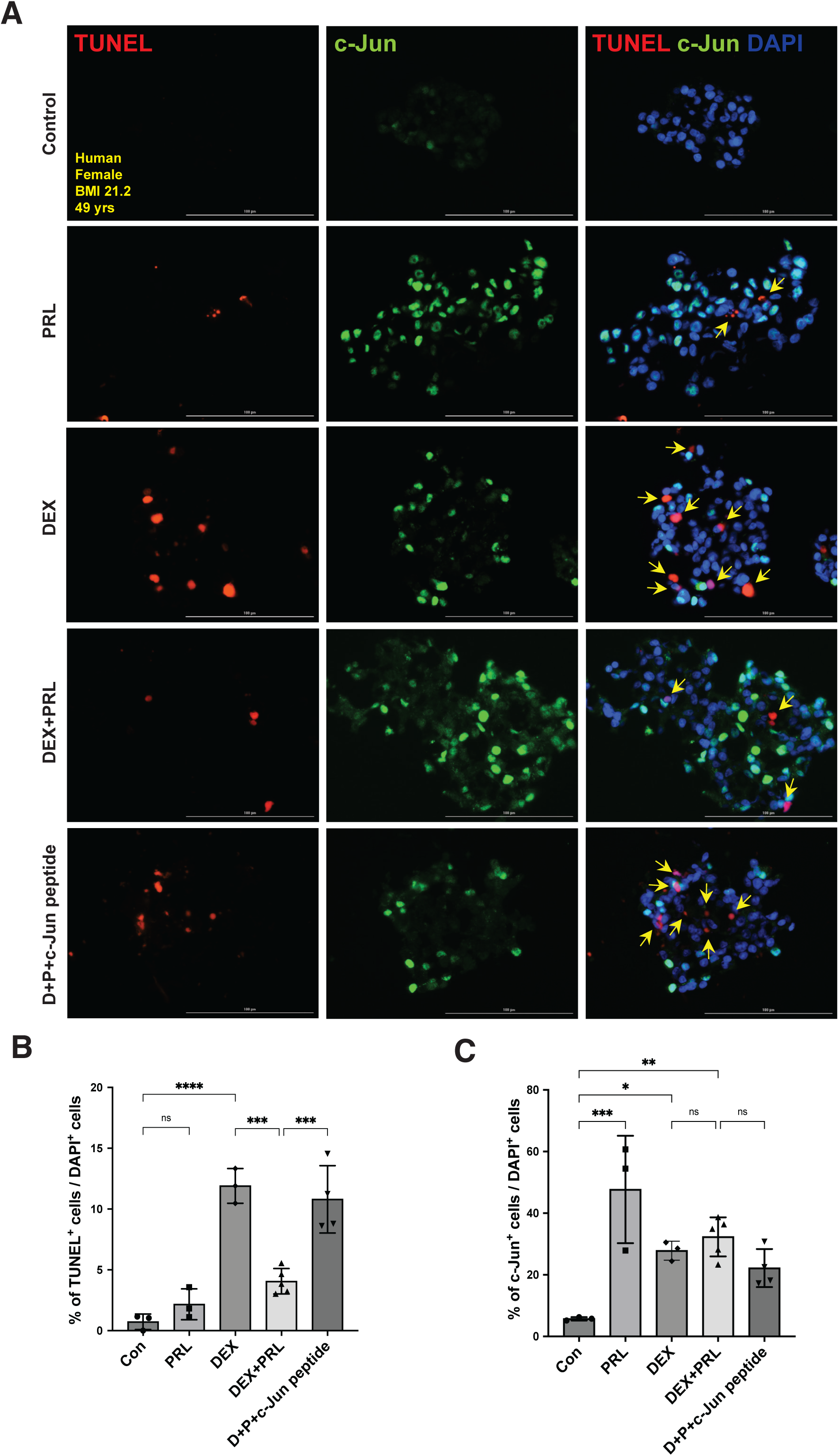
Human islets are also protected against induced apoptosis via c-Jun. (A) Immunostaining of cultured human islets for c-Jun (green) following treatment with PRL, DEX, DEX+PRL, or DEX+PRL+c-Jun peptide (inhibitor). Apoptosis is indicated by TUNEL+ nuclei (red). All nuclei are stained with DAPI (blue). (B) Quantification of percentage of apoptosis (TUNEL+ cells) / total islet cells (DAPI+) (C) Quantification of Jun+ cells / total islet cells (DAPI+). Data in B and C were analyzed using a 2-way ANOVA. Data are represented as mean ± SEM. Statistical significance was defined as *p ≤ 0.05, **p ≤ 0.01, ***p ≤ 0.001, and ****p ≤ 0.0001.

### 2.4 c-Jun knockdown in MIN6 cells blocks the protective effect of PRL on DEX-induced apoptosis

Islets are comprised of multiple cell types, and while only murine β-cells express PRLR at detectable levels, it remained possible that non-cell autonomous effects or secondary effects were responsible for c-Jun-mediated survival of β-cells following PRL treatment of cultured islets. To establish c-Jun function within β-cells is critical for protection against DEX-induced apoptosis, we examined the consequences of c-Jun loss in the MIN6 cell culture model of murine β-cells [31] using siRNA knockdown. Basal levels of c-Jun mRNA and protein were very low in MIN6 cells but were induced robustly by PRL, whereas siRNA-mediated knockdown of c-Jun prevented the c-Jun mRNA and protein increase following PRL treatment (**Figure 6A**). Using siRNA-mediated knockdown of *Jun*, we next examined the functional consequences of c-Jun loss in MIN6 cells. PRL treatment increased MIN6 viability as measured by the MTT assay (**Figure 6B**). Treatment with DEX substantially reduced MIN6 cell viability, which was partially rescued by PRL treatment. However, PRL failed to improve viability when either pharmacologic (c-Jun peptide), or genetic silencing (*Jun* siRNA) methods were used to inhibit c-Jun. Together, these results indicate that c-Jun protein and activity are required for PRL-mediated protection against DEX-induced apoptosis in MIN6 cells as in primary β-cells and are not due to interactions with other islet cell types.

**Figure 6.**
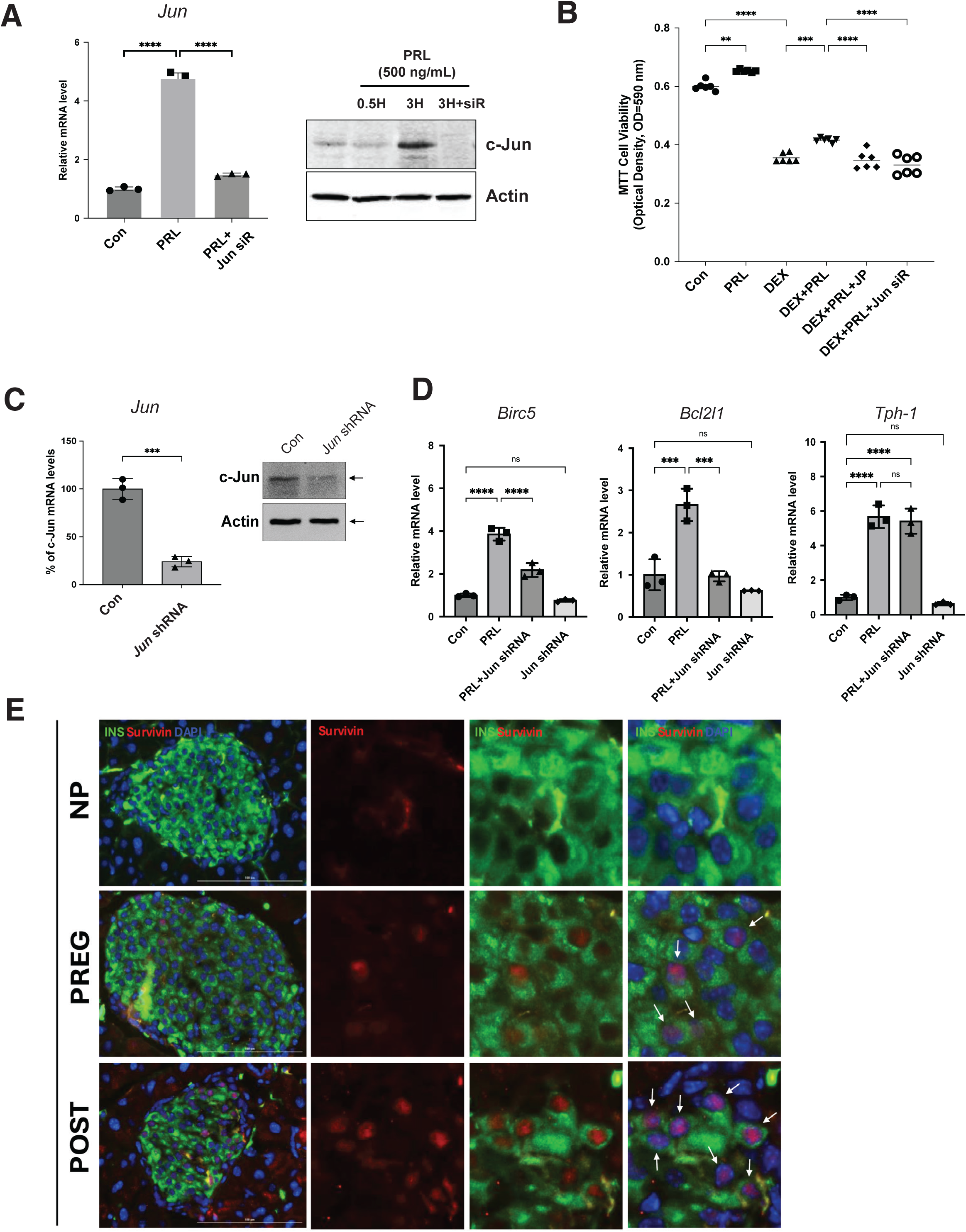
*Jun* knockdown in MIN6 cells impairs cell survival and induction of pro-survival genes. (A) Knockdown of *Jun* by siRNA transient transfection in MIN6 cells. c-Jun mRNA and protein levels were confirmed by Q-PCR and western blot after PRL treatment. (B) MTT assay for survival of MIN6 cells treated with PRL, DEX, with or without c-Jun peptide or siRNA (knockdown) of *Jun*. (C) c-Jun mRNA and protein levels were confirmed by Q-PCR and western blot after lentiviral transduction in MIN6 cells. (D) Expression of c-Jun targets in MIN6 cells after *Jun* knockdown by lentiviral transduction and PRL treatment was quantified using Q-PCR (E) Survivin immunostaining of sections from murine pancreas of non-pregnant (NP), pregnant (PREG) and postpartum (POST). Insulin (green) and survivin (red); nuclei marker with DAPI (blue). White arrows indicate survivin positive β-cells in both PREG and POST but none in NP mice. Statistical testing was performed with a 2-way ANOVA in (A), (B) and (D) and with student’s t-text in (C). Data are represented as mean ± SEM. Statistical significance was defined as **p ≤ 0.01, ***p ≤ 0.001, and ****p ≤ 0.0001.

### 2.5 Identification of downstream mediators of c-Jun anti-apoptotic effects in β-cells

To investigate downstream effectors by which Jun might promote β-cells survival following pro-apoptotic stress, we investigated candidate genes known to act as anti-apoptotic factors and regulated during pregnancy or postpartum within islets. Survivin, a member of the inhibitor of apoptosis family and encoded by the *Birc5* gene is gestationally regulated by PRLR signaling in β-cells [11, 32]. To ensure robust and consistent knockdown of *Jun*, we used lentiviral transduction of MIN6 cells with shRNA against *Jun* for gene expression studies. Lenti-shRNA-*Jun* treatment of MIN6 cells significantly reduced both mRNA and protein levels of c-Jun (**Figure 6C**). Next, we examined the effects of c-Jun knockdown by sh-RNA on expression of *Birc5*, as well as another anti-apoptotic gene *Bcl2l1* (Bcl-xL) which is also induced by PRL and promotes β-cell survival [29]. PRL treatment upregulated *Birc5* in MIN6 Cells; this was significantly attenuated by c-Jun knockdown (**Figure 6D**). This was specifically attributed to Jun knockdown as JunB and JunD were not blunted by Jun shRNA (**Supplementary Figure 5**). *Bcl2l1* was induced by PRL, but the induction was completely abrogated by c-Jun knockdown. By contrast, induction of another PRL/PRLR target gene, *Tph1,* which regulates islet serotonin production (**Figure 6D**) was not affected by *Jun* expression. Jun knockFurthermore, we confirmed survivin protein was increased in vivo during pregnancy and postpartum by immunostaining (**Figure 6E**). Together, these results indicate that anti-apoptotic factors survivin and Bcl-xL are both regulated by c-Jun, providing a mechanistic connection between c-Jun and established mediators of β-cell survival.

### 2.6 c-Jun in human pregnancy and gestational diabetes

We sought to determine whether the regulation of c-Jun during gestation was relevant to human biology. We obtained sections of pancreata from pregnant females and non-pregnant controls from the Network of Pancreatic Organ Donors (nPOD) (**Table 1**). Non-pregnant donors with similar ages and BMIs exhibited no c-Jun immunoreactivity within either the islets or exocrine pancreas. Human donors from pregnancy both exhibited significantly increased c-Jun levels in islet β-cells, but not α-cells (**Figure 7A&B**). Incidentally we observed, c-Jun immunoreactivity within the exocrine pancreas in both pregnancy donors. However, in human pregnancy complicated by GDM, c-Jun induction in islets was absent (**Figure 7C-D**). The detection of c-Jun staining in GDM exocrine tissue indicates this decrease was specific to β-cells and not generalized loss of c-Jun with GDM.

**Figure 7.**
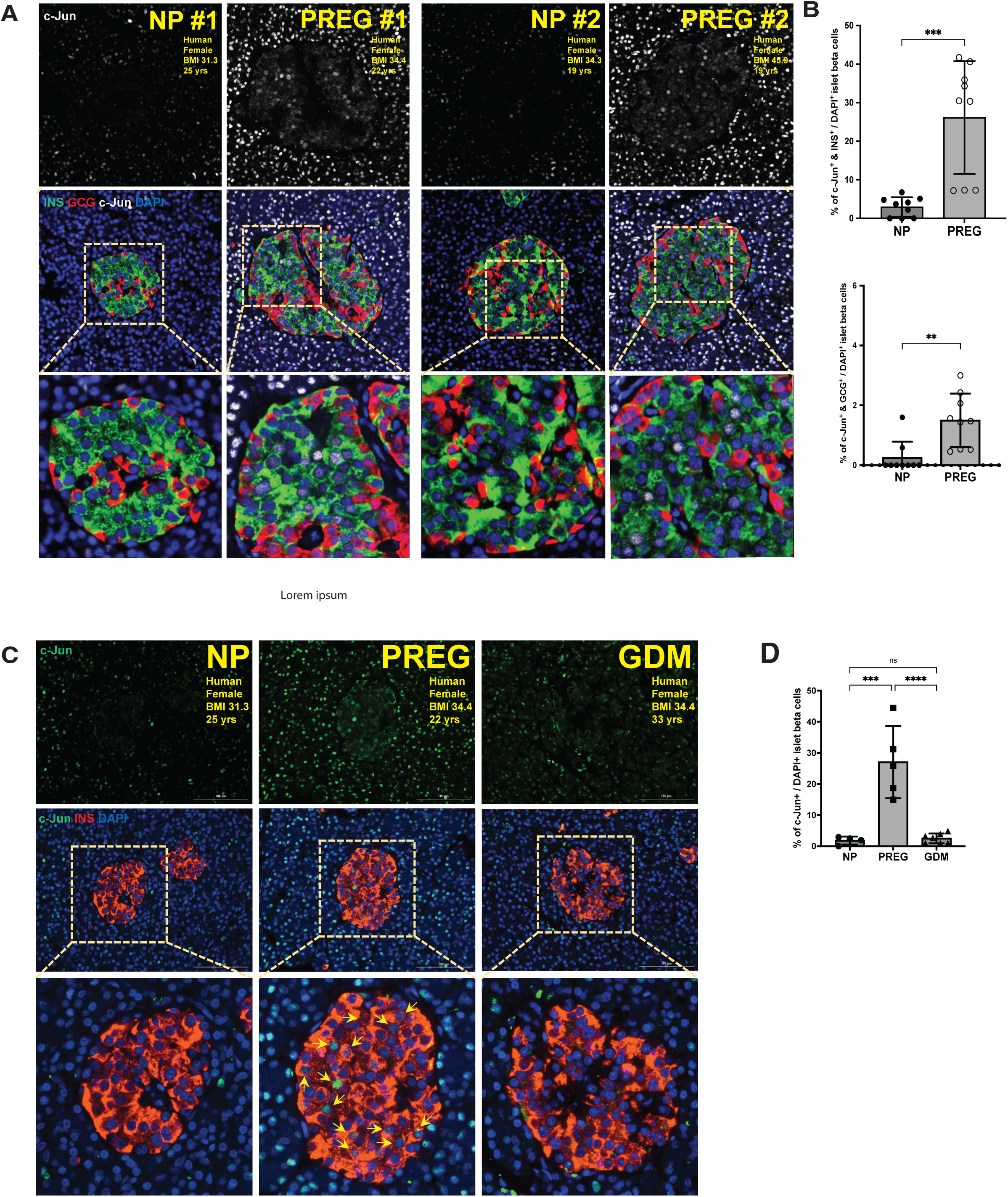
c-Jun induction is conserved in human pregnancy but absent in pregnancy complicated by gestational diabetes. (A) Immunostaining of sections from human pancreas non-pregnant and pregnant donors. Insulin (green), glucagon (red), and c-Jun expression (white); nuclei marker with DAPI (blue). (B) Quantification of c-Jun+ nuclei in β-cells (Ins+) or alpha-cells (Gcg+) as a percentage of islet nuclei (DAPI). (C) Immunostaining of sections from human pancreas non-pregnant, pregnant, and pregnant with GDM donors. c-Jun (green), insulin (red), and nuclei marker with DAPI (blue). (D) Quantification of c-Jun+ nuclei in β-cells (Ins+) as a percentage of total islet cells (DAPI+). Data in (B) was analyzed using a student’s t-test and in (D) using a 2-way ANOVA. Data are represented as mean ± SEM. Statistical significance was defined as **p ≤ 0.01, ***p ≤ 0.001, and ****p ≤ 0.0001.

Together, these studies indicate that gestational regulation of c-Jun is within β-cells is conserved in humans and mice, despite species differences in gestational biology. The loss of c-Jun in human pregnancy with GDM suggests that loss of c-Jun may contribute to GDM pathogenesis.

## 3. Discussion

In this study, we showed that prolactin signaling in late gestation and postpartum periods induces c-Jun expression via MAPK signaling. In turn, c-Jun activation protects β-cells against DEX-induced apoptosis by increasing expression of survivin and Bcl-xL pro-survival factors, thereby contributing to β-cell survival during postpartum regression. Intriguingly, this adaptation is absent in human pregnancies complicated by GDM.

Survivin and Bcl-xL have been previously implicated as critical for β-cell survival in other contexts. Survivin promotes survival of β-cells in a transplantation model [33]. Similarly, Bcl-xL also regulates β-cell survival vs. apoptosis in different settings [34]. Prior studies showed Bcl-xL is induction by lactogens in both mouse islets [35] and INS-1 cells [36] where it protects against apoptosis. Thus, consistent with their role in other contexts, our findings demonstrate expression of anti-apoptotic genes is critical to the survival of β-cells during postpartum regression. In contrast to survivin, Bcl-xL induction is mediated by Stat5/Jak2 signaling [29, 37]. Most likely, the protective effects of prolactin are mediated by multiple signal transduction pathways. This should motivate studies to investigate whether MAPK and Stat5/Jak2 activation promotes β-cell survival and thereby maintains β-cell mass under conditions that increase β-cell stress and cell death.

Interestingly, survivin is also necessary for β-cell proliferation during early stages of pregnancy [32, 38]. This bifunctional role for survivin as both apoptosis inhibitor and pro-proliferative factor has been reported in cancer biology [39]. Examining precisely how survivin protects postpartum β-cells from apoptosis, and whether it contributes to β-cell proliferation in the postpartum lactation period [19] should be determined in future studies. Furthermore, while both survivin and Bcl-xL were logical candidates to examine as effectors of c-Jun’s pro-survival effects, a comprehensive and unbiased identification of genome-wide c-Jun targets within postpartum β-cells will be helpful in expanding our understanding of how c-Jun regulates pro-survival and other β-cell responses.

We showed that β-cell induction of c-Jun during gestation is conserved in mouse and humans. The reduction of c-Jun expression in human pregnancies complicated by GDM is intriguing, as it suggests a potential mechanism of increased susceptibility to β-cell apoptosis and loss following pregnancy. Clinical studies have established that women with either GDM or glucose intolerance (GIGT) during pregnancy have a decline in β-cell function during the first year postpartum [40] and maternal prolactin inversely correlates with risk of postpartum diabetes [41]. If so, this could provide a potential mechanistic connection between GDM, increased β-cell loss postpartum, and the future risk of T2D in mothers with lost or insufficient c-Jun expression. Additional work is needed to ascertain whether GDM pregnancies have increased β-cell apoptosis in vivo during the postpartum period. Formal demonstration that c-Jun regulates β-cell mass in vivo requires generation of conditional β-cell loss of function animal models that would be expected to exhibit greater postpartum loss of β-cells with the absence of c-Jun. Finally, is the absence of c-Jun in human GDM pregnancy indicative of functional defects within the β-cell caused by hyperglycemia, or perhaps indicates a broader failure of downstream PRLR signaling in human GDM that represents “prolactin resistance”? While human gestational and postpartum islet studies are challenging because of a lack of samples, identification of c-Jun provides a specific target and hypotheses to interrogate.

One weakness of this study is that we focused on a single member of the c-Jun family. Our prior data showed gestational and postpartum changes in expression of other Jun family members *JunB* and *JunD*, as well as *Fos* family members [7]. Emerging evidence suggests that individual Jun family members exhibit unique specificity to regulate cellular functions such as in cancer [42], infection/inflammation and neuronal development and metabolism [20]. Thus, it is likely that further studies to delineate specific roles for individual Jun family members in regulating β-cell function will be illuminating. For example, a recent study indicating that JunD maladaptively contributes to oxidative stress and apoptosis in β-cells following glucolipotoxic stress [43] contrasts with our finding that c-Jun is regulated by gestational rather than nutritional stress.

Our studies lead us to speculate that modulating c-Jun activity or, more broadly, pathways downstream of PRLR signaling could provide a therapeutic strategy to maintain or promote functional β-cell mass in the treatment of gestational diabetes. Finally, recent transcriptomic studies identified *Jun* and *Fos* amongst “nutrient-responsive” transcription factors driving early postnatal developmental proliferation and expansion of β-cells [44]; immediate early gene transcription factors (IEG-TFs) including *Fos* and *Nr4a1* may contribute to β-cell adaptations following acute switching from the fasting to feeding state [45]. Together with our findings that Jun, Fos, and other IEG-TFs are induced within gestational or postpartum β-cells [7], these intriguing studies suggest that IEG-TFs play a wider role in regulating β-cell metabolism, and warrant further investigation.

## 4. Methods

### 4.1 Animals

All experimental protocols and animal care were reviewed and approved by the Johns Hopkins University Animal Care and Use Committee. Mice were fed regular chow diet and maintained in rooms with 12-hour light/dark cycle and constant temperature and humidity. All mice had free access to food and water. 8-week-old virgin C57Bl/6J females were mated with B6 males then single-housed after vaginal plugs were visualized (Jackson Labs, #000664). Gestational day (GD) was determined by presence of vaginal plugs (0.5). Islets were isolated from age-matched virgin females, pregnant females at GD16.5 and postpartum females at day 5 following parturition (postpartum day 0). Male mice were provided high-fat, high-sucrose diet (HFHS) consisting of 58% Kcal fat and 13% Kcal sucrose for 24 weeks (Research Diets, Cat# D09071702).

### 4.2 Human and mouse islet procurement and culture conditions

Mouse islet isolation was performed using collagenase digestion of the pancreas and density centrifugation as previously described [7]. Murine islets were then handpicked in HBSS buffer containing 2% fetal bovine serum (FBS, Cat # 26140079, Thermo Fisher Scientific (TFS), Waltham, MA) using a dissecting microscope. Human islets were provided by the Integrated Islet Distribution Program (IIDP); see **Table 1**. Culture media used for mouse and human islets was RPMI1640 (Cat # 22400089, TFS) supplemented with 10% FBS and 25 mM HEPES (Cat # 15630080, TFS).

### 4.3 Immunofluorescence (IF) & Imaging analyses

Human Pancreatic tissues were provided by the Network for Pancreatic Organ Donors with Diabetes (nPOD); see **Table 1**. Mouse pancreata were fixed for 30 min at room temperature in 4% paraformaldehyde (PFA, Cat # 100503-917, VWR, Radnor, PA). Fixed tissues were incubated sequentially in 10, 20, and 30% sucrose for 1h each, then embedded in OCT compound (Cat # 23-730-571, Fisher Scientific, Hampton, NH). Human and mouse islets treated with PRL or other drug combinations were embedded in agarose-OCT compound, and they were sectioned to 5 μm thickness by cryostat. For immunofluorescence staining, paraffin-embedded human pancreas tissues were deparaffinized, hydrated and antigen retrieval performed using sodium citrate buffer (10 mM, pH 6.0) in a pressure cooker. Samples were washed with PBS then permeabilized with 0.2% containing Triton X-100 (Cat # T9284, Millipore Sigma). After washing, samples were incubated with the blocking buffer (3% BSA, Cat # A9647, Millipore Sigma and 5% normal goat serum, Cat # 5425, Cell Signaling Technologies (CST) in PBS) in a humidified chamber for 1 h at room temperature. The primary antibodies: anti-insulin (1:500, polyclonal guinea pig, Cat # IR00261-2, Agilent, Santa Clara, CA), anti-glucagon (1:500, monoclonal mouse, Cat # G2654, Millipore Sigma), anti c-Jun (1:200, Cat # 9165, CST) were incubated overnight at 4 °C. Secondary antibodies were diluted to 1:1000, then incubated at room temperature for 1 h (anti-guinea pig, anti-mouse, or anti-rabbit IgG antibodies conjugated with Alexa Fluro 488, Cat # TA11008, 546, Cat # A21123, or 647 Cat # A21244 all from TFS). Samples were mounted using Vectashield mounting medium containing DAPI (Cat # H-1200, Vector Laboratories, Newark, CA). The TUNEL assay was conducted with the Click-iT Plus TUNEL Alexa Fluor Imaging Assay kit (Cat # C10617, TFS) per manufacturer’s instructions. Images were captured using a Lionheart FX automated cell imaging microscope (Agilent-BioTek, Santa Clara, CA) and LSM700 inverted laser scanning confocal microscope (Carl Zeiss, Thornwood, NY).

### 4.4 RNA Extraction and Reverse Transcription-Quantitative PCR (Q-PCR)

Total RNA isolation and cDNA synthesis, Real-time Q-PCR as previously described [7]. *Rps29* was used as a reference gene. See **Supplementary Table 1** for details regarding primers.

### 4.5 MIN6 cells

MIN6 cells, a mouse pancreatic β-cell line derived from insulinomas in transgenic mice (originating in [46]) were provided by Maria Golson (Johns Hopkins School of Medicine, Baltimore, MD). MIN6 cells were cultured in Dulbecco’s Modified Eagle’s Medium (DMEM, Cat # 30-2002, ATCC, Manassas, VA) containing 15% FBS (Cat # 26140079, Thermo Fisher Scientific (TFS)), 10mM HEPES (Cat # 15630080, TFS), 100 units penicillin -streptomycin (Cat # 15140122, TFS) and 70 μM β-Mercaptoethanol (Cat # 21985023, TFS). Cells were incubated in a humidified incubator at 37°C with 5% CO_2_.

### 4.6 Western blot

Freshly isolated mouse islets or MIN6 cells after combination of treatments were incubated in RIPA lysis buffer with protease and phosphatase inhibitor cocktail EDTA-free (Cat # A32961, TFS) for 45 min on ice. The lysed soluble proteins after centrifugation at 12,000 rpm for 15 min at 4 °C were collected, and the protein concentration was determined by a BCA protein assay kit (Cat # 470203834, TFS). Equal amounts of proteins (25 μg) for each sample were subjected to 8-15% sodium dodecyl sulfate-polyacrylamide gel electrophoresis (SDS-PAGE) and transferred onto nitrocellulose membranes. The membranes were then blocked with the OneBlock Western blocking solution (Cat # 10-313, Genesee Scientific) at RT for 1 h and the membranes were incubated with the primary antibody diluted in blocking solution. After washing the membranes, secondary antibody reactions were conducted with an appropriate source of antibody conjugated with infrared fluorescent dyes (Rabbit Cat # 926-32211 and Mouse Cat # 926-32210, Li-Cor, Lincoln, NE). The signals were detected in the Odyssey Imaging System (Li-Cor, Lincoln, NE). The intensity of some band was quantified using the ImageJ software (http://rsb.info.nih.gov/ij/; National Institutes of Health, Bethesda, MD). See **Supplementary Table 2** for details regarding the primary antibodies.

### 4.7 Cell viability assay (MTT, 3-(4,5-Dimethylthiazol-2-yl)-2,5-Diphenyltetrazolium Bromide)

To shed further light on the effect of c-Jun on cell viability, we conducted MTT cell viability assay. MIN6 cells were seeded in 6-well culture plates at a density of 1 × 10^5^ cells per well and cells grown until they attained 50% confluency. Cells were treated with **c-Jun peptide** (Cat # 1989, Bio-Techne/Tocris, blocks JNK/c-Jun interaction and c-Jun phosphorylation [47]) in advance and additional incubation was performed with PRL or combination with dexamethasone (DEX) for 24 h in the presence of c-Jun peptide. After 24 h incubation, culture medium was discarded and replaced with a fresh medium containing 0.5 mg of MTT (Cat # M6494, TFS) dissolved in PBS (pH=7.2). After further 3 h incubation, the formazan purple crystals were dissolved with dimethyl sulfoxide (DMSO, Cat # D2650, Millipore Sigma). Dissolved crystals were transferred to 96 well plate and the intensity of the color in each well was measured at a wavelength of 590 nm using a microplate spectrophotometer reader (Agilent Bio-Tek, Epoch, Winooski, VT).

### 4.8 Chemicals and Inhibitors used in islet and MIN6 studies

To determine the downstream signaling of PRL and PRLR for c-Jun upregulation in β-cell, AG490 (Stat5 inhibitor, Cat # 0414, Bio-Techne/Tocris, Minneapolis, MN), LY294002 (PI3K inhibitor, Cat # 1130, Bio-Techne/Tocris), U0126 (MEK1/2 (MAPK), inhibitor Cat # 1144, Bio-Techne/Tocris), and Cycloheximide (protein synthesis inhibitor, Cat # 0970, Bio-Techne/Tocris) were used in isolated murine and human islets. Briefly, islets were incubated with each inhibitor for 30 min respectively prior to PRL treatment then cultured with PRL in the presence of each inhibitor for additional 24 h. After incubation, islets were collected for protein extraction to perform Western blotting.

### 4.9 Knockdown of *Jun* in MIN6 cells

To knock down *Jun*, both siRNA transient transfection and lentivirus transduction with sh-RNA against the *Jun* gene were used. 100 nM of *Jun* siRNA (Cat # 6203, CST) was introduced into MIN6 cells using Lipofectamine RNAiMAX transfection reagent (Cat # 13778100, TFS) in Opti-MEM medium (Cat # A4124801, TFS), according to the manufacturer’s instructions and cells were incubated for 48 h in the presence of *Jun* siRNA. A lentivirus containing green fluorescence protein (GFP) with a modified pLKO.1 vector and optimized packaging construct ratios (Millipore Sigma, MISSION shRNA, SHCLNV, Cat # TRCN0000229528) was used to create a stable gene knockdown for *Jun* in MIN6 cells. To enhance lentivirus transduction, hexadimethrine bromide (Cat # H9268, Millipore Sigma) was added to the medium to a final concentration of 8 μg/mL. 7.5×10^3^ lentiviral particles were added to MIN6 cells cultured in 12-well plates and incubated for 24 h. After incubation with lenti-shRNA-*Jun or* lenti*-GFP* controls, medium was changed and cells were incubated for an additional 48 h.

### 4.10 Quantification and statistical analysis

Statistical and graphing analyses were performed using GraphPad Prism 10.0 software (GraphPad Software Inc., La Jolla, CA) and Microsoft Excel (for mac v16.87). All data are expressed as means ± SEM with p< 0.05 considered statistically significant unless otherwise noted. Statistical significance was denoted within figures as *p ≤ 0.05, **p ≤ 0.01, ***p ≤ 0.001, and ****p ≤ 0.0001.

## Author Contributions

R.R.B. conceptualized the study and designed the experiments. J-Y. C. and N. R-O performed experiments and analyses. R.R.B wrote the manuscript. All authors read, edited, and approved the manuscript. R.R.B obtained funding and supervised and administered the project.

## Acknowledgements

This work was supported by a National Institutes of Health (NIH) grant R01-DK120761 (to R.R.B). N.R-O is supported by an American Diabetes Association (ADA) Postdoctoral Fellowship (11-23-PDF-34).

## Data Availability

Data will be made available on request.

## Supplementary Data

Supplementary Data are included as a separate file.

---

**Supplementary Table 1:** Q-PCR Primers.

| Oligonucleotide | Source | Identifier |
| --- | --- | --- |
| *Tph-1* | Thermo Fisher Scientific | Mm.PT.58.8815029 |
| *Rps29* | Thermo Fisher Scientific | Mm.PT.58.21577577 |
| *c-Jun* | Integrated DNA Technologies | Mm.PT.58.32691984.g |
| *JunB* | Integrated DNA Technologies | Mm.PT.58.30360909.g |
| *JunD* | Integrated DNA Technologies | Mm.PT.58.32598468.g |
| *Birc5* | Thermo Fisher Scientific | Mm00599749_m1 |
| *Bcl2l1* | Thermo Fisher Scientific | Mm00437783_m1 |

**Supplementary Table 2:** Primary antibody.

| Antibodies | Source | Identifier |
| --- | --- | --- |
| c-Jun | Cell Signaling Technology | Cat # 9165, RRID:AB_2130165 |
| phospho-c-Jun | Cell Signaling Technology | Cat # 9164, RRID:AB_330892 |
| Stat5 | Cell Signaling Technology | Cat # 94205, RRID:AB_2737403 |
| p-Stat5 | Cell Signaling Technology | Cat # 9351, RRID:AB_2315225 |
| Erk1/2 | Cell Signaling Technology | Cat # 9102, RRID:AB_330744 |
| p-Erk1/2 | Cell Signaling Technology | Cat # 9101, RRID:AB_331646 |
| Actin | Cell Signaling Technology | Cat # 4970, RRID:AB_2223172 |
| GAPDH | Santa Cruz Biotechnologies | Cat # sc-47724, RRID:AB_627678 |
| Survivin | Santa Cruz Biotechnologies | Cat # sc-17779, RRID:AB_628302 |
| Glucagon | Millipore Sigma | Cat # G2654, RRID:AB_259852 |
| Insulin | Agilent | Cat # IR00261-2 |

