## Supplemental Figures for "c-Jun regulates postpartum β-cell apoptosis and survival downstream of prolactin signaling"

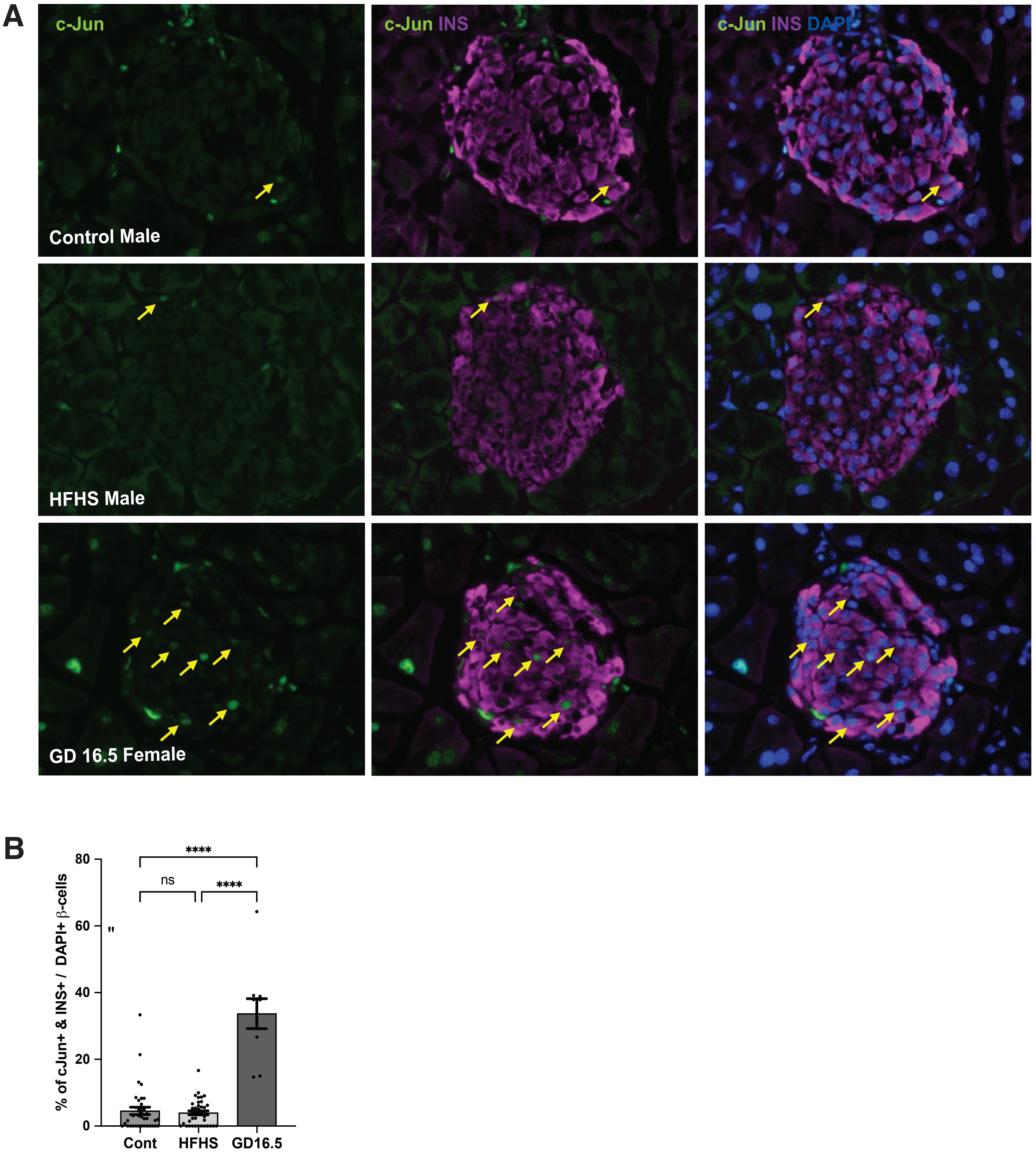


**Supplementary Figure S1. Unlike pregnancy, dietary stress does not induce c-Jun.**

**(A)** Male mice fed a high-fat, high-sucrose diet do not exhibit increased c-Jun immunoreactivity. GD16.5 female mice serve as a positive control for c-Jun staining within β-cells.

**(B)** Quantitation of **(A)** indicating percentage of β-cells that are c-Jun positive. (c-Jun+Ins+ cells / Ins+DAPI+ cells).

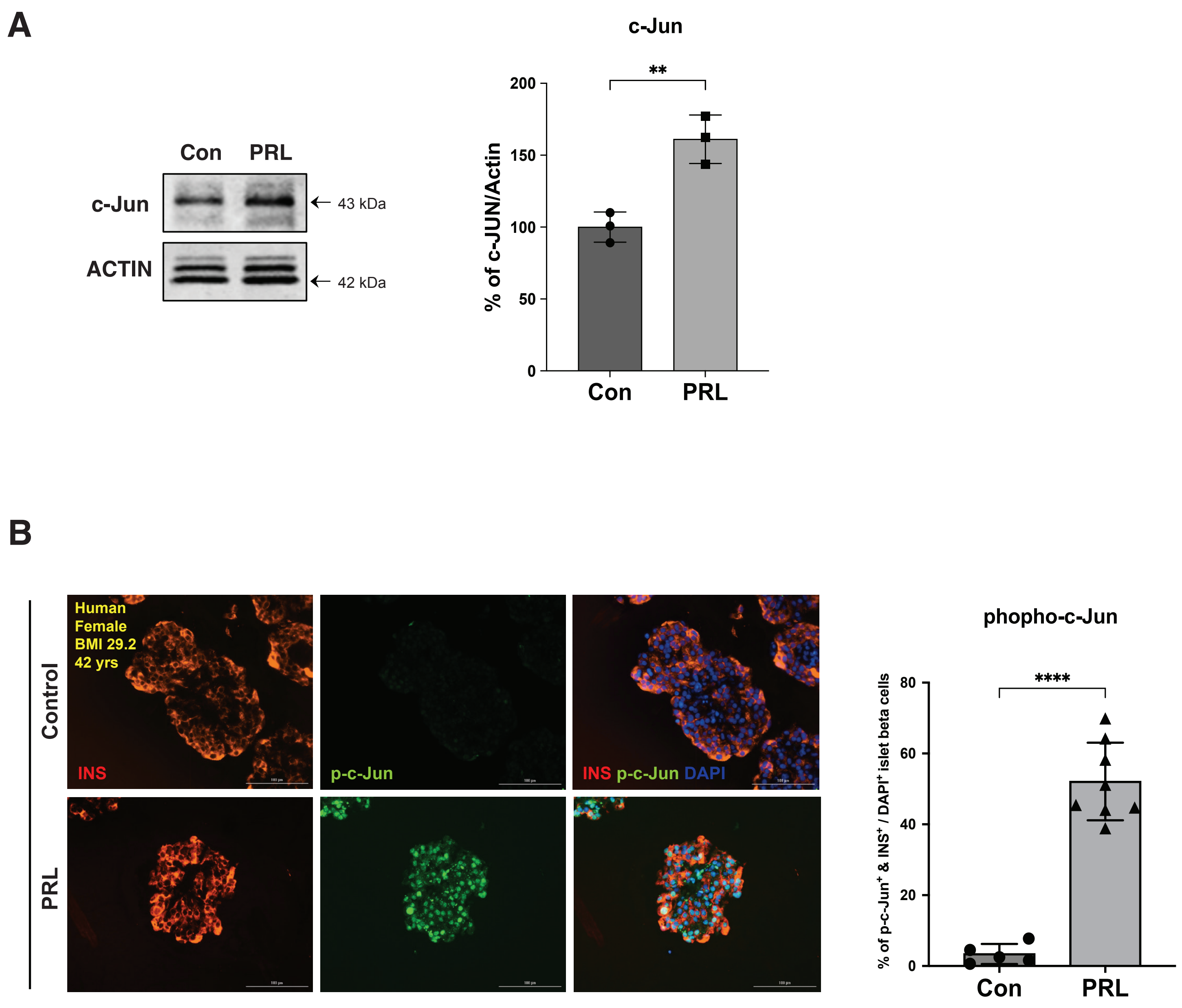


**Supplementary Figure S2. c-Jun in isolated islets treated with prolactin.**
**(A)** Western blot of islets from female mice treated with prolactin for 24 hours show increased c-Jun.
**(B)** Immunostaining of human female islets treated with recombinant human prolactin show increase in phospho-c-Jun in β-cells**.**

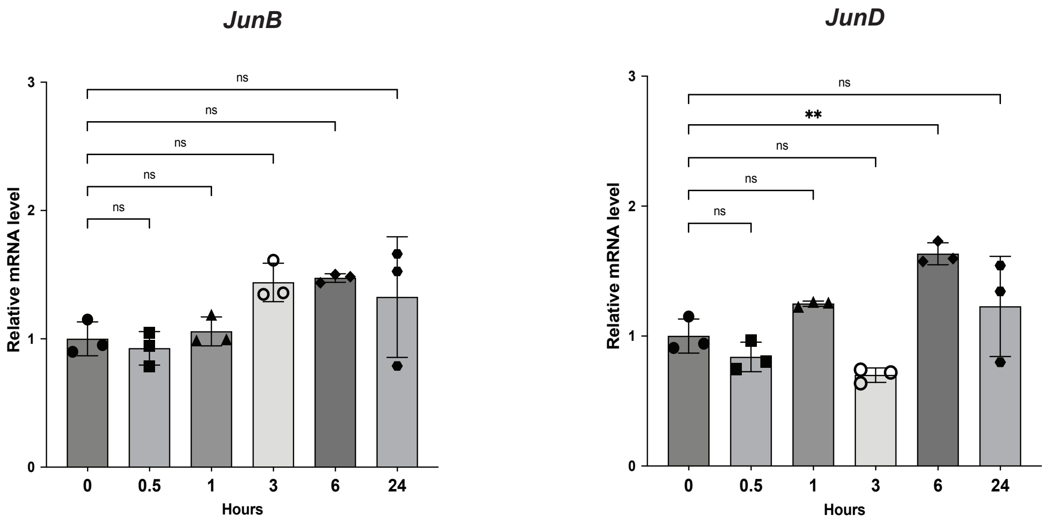


**Supplementary Figure S3. c-Jun family members JunB and JunD are not increased by prolactin treatment**. By contrast with *c-Jun*, *JunB* and *JunD* mRNA are not induced follow treatment of murine islets with recombinant mouse prolactin.

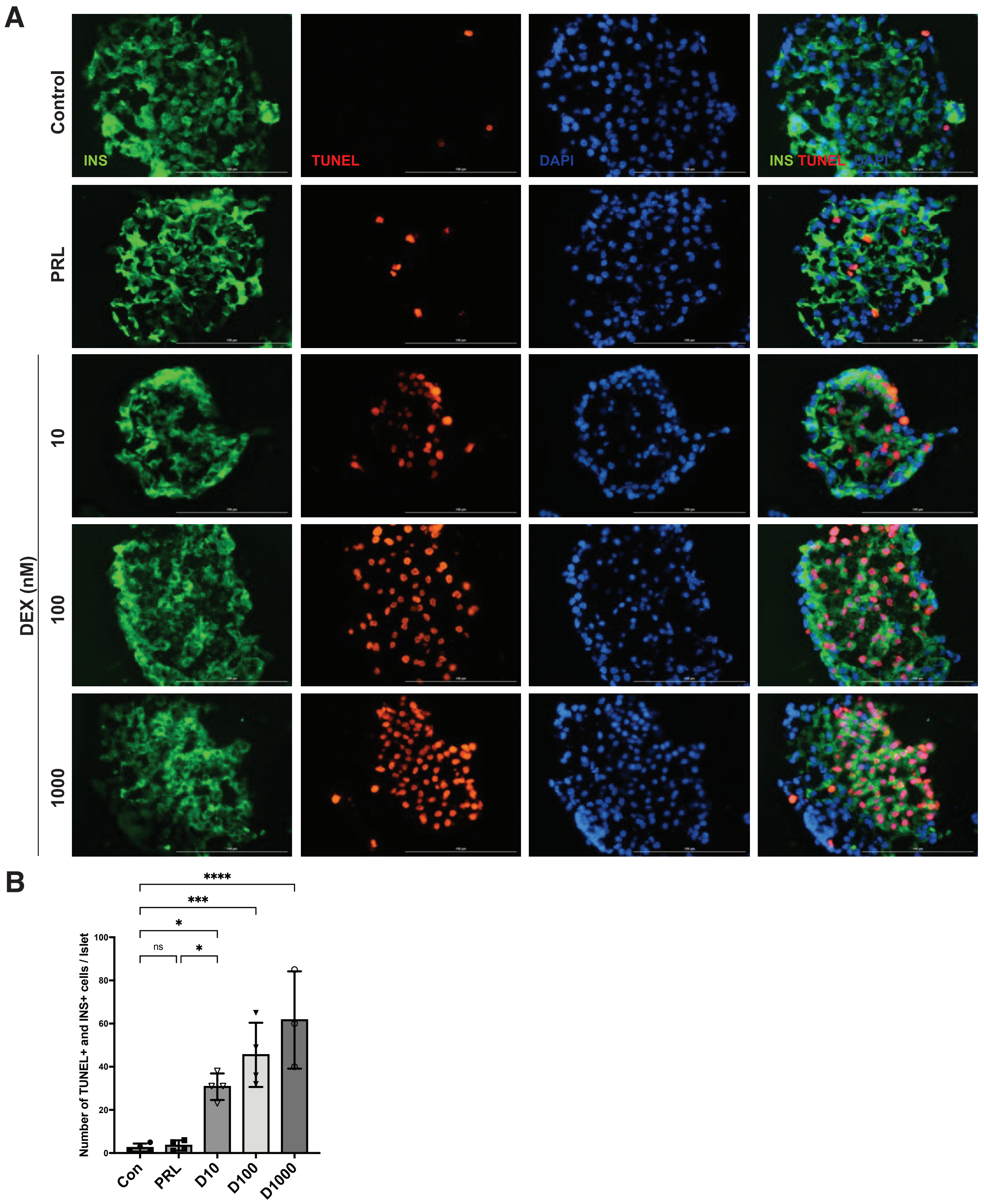


**Supplementary Figure S4. Dexamethasone dose response for β-cell apoptosis.**

(**A**) Immunofluorescense of islets with Insulin (green), TUNEL (red) and DAPI (blue) staining.

(**B**) Quantification of TUNEL+ β-cells per islet.

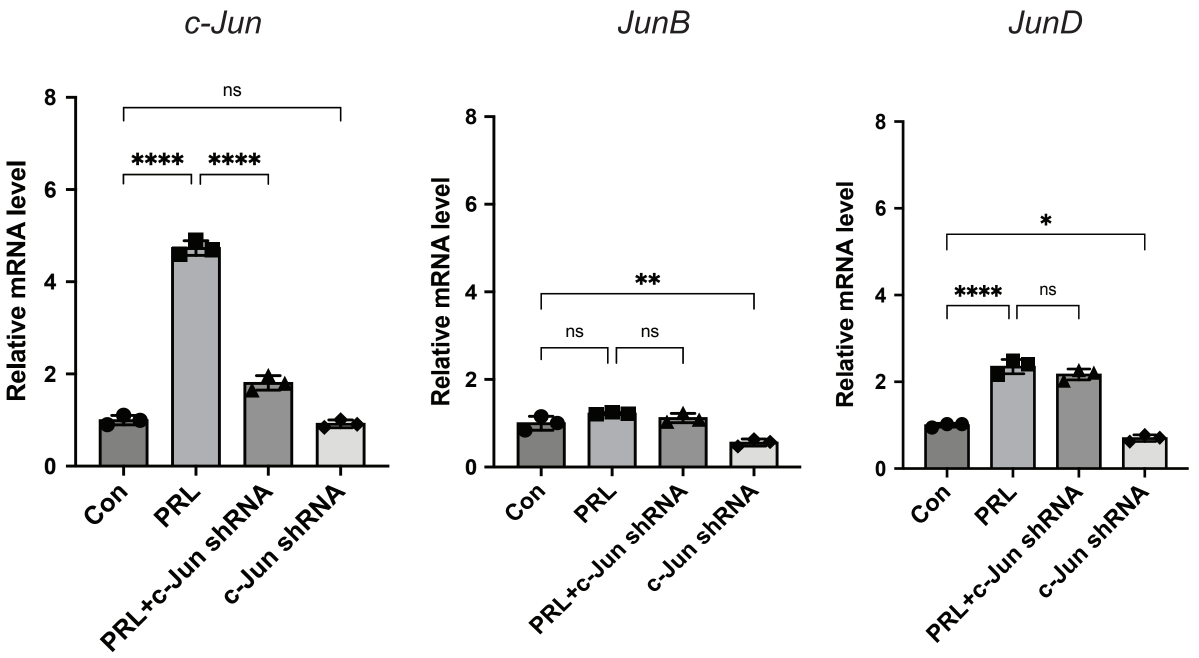


**Supplementary Figure S5. MIN6 knockdown of c-Jun using shRNA.** Expression of *c-Jun, JunB* and *JunD* genes following c-Jun knockdown with lentivirus transduction. MIN6 cells were cultured with lentivirus for 24 h and replaced with new medium. Cells were then cultured for an additional 48 h, then treated with PRL for 24 h. Induction of *c-Jun* following PRL is blunted by Jun-shRNA unlike *JunB* and *JunD*.
